# TCGAnalyzeR: a web application for integrative visualization of molecular and clinical data of cancer patients for cohort and associated gene discovery

**DOI:** 10.1101/2023.01.20.524925

**Authors:** Talip Zengin, Başak Abak Masud, Tuğba Önal-Süzek

**Affiliations:** Department of Molecular Biology and Genetics, Mugla Sitki Kocman University, Mugla, 48000, Turkiye; Department of Bioinformatics, Mugla Sitki Kocman University, Mugla, 48000, Turkiye; Department of Computer Engineering, Mugla Sitki Kocman University, Mugla, 48000, Turkiye

## Abstract

**Motivation:** The vast size and complexity of The Cancer Genome Atlas (TCGA) database with multidimensional molecular and clinical data of ~11,000 cancer patients of 33 cancer types challenge the effective utilization of this valuable resource. Therefore, we built a web application named TCGAnalyzeR with the main idea of presenting an integrative visualization of mutations, transcriptome profile, copy number variation and clinical data allowing researchers to facilitate the identification of customized patient cohorts and gene sets for better decision-making for oncologists and cancer researchers.

**Results:** We present TCGAnalyzeR for integrative visualization of pre-analyzed TCGA data with the several novel modules: (i) Simple nucleotide variations with driver prediction; (ii) Recurrent copy number alterations; (iii) Differential expression in tumor versus normal, with pathway enrichment and the survival analysis; (iii) TCGA clinical data and survival analysis; (iv) External subcohorts from literature, curatedTCGAData and BiocOncoTK R packages; (v) Internal patient clusters determined using iClusterPlus R package or signature-based expression analysis. TCGAnalyzeR provides clinical oncologists and cancer researchers interactive and integrative representations of these multi-omic, pan-cancer TCGA data with availability of subcohort analysis and visualization. TCGAnalyzeR can be used to create their own custom gene sets for pan-cancer comparisons, to create custom patient subcohorts comparing external subcohorts (MSI, Immune, PAM50, Triple Negative, IDH1, miRNA, etc) along with our internal patient clusters, to visualize cohort-centric or gene-centric results along with pathway enrichment and survival analysis graphically on an interactive web tool.

**Availability:** TCGAnalyzeR is freely available on the web at http://tcganalyzer.mu.edu.tr.

**Supplementary information:** Supplementary data are available at *Bioinformatics* online.

## 1 Introduction

The sheer scale and complexity of The Cancer Genome Atlas (TCGA) data (The Cancer Genome Atlas Research Network. *et al.*, 2013) offers great potential for scientific discovery, but the challenges to effective use of this valuable resource by biologists and clinicians have led to the development of several visualization tools such as cBioPortal (Cerami *et al.*, 2012; Gao *et al.*, 2013), Firebrowse (Deng *et al.*, 2017), and University of California, Santa Cruz (UCSC) Xena (Goldman *et al.*, 2020). Among these tools, cBioPortal is the most preferred due to its interactive exploration of larger and up-to-date cancer datasets. OncoKB (Chakravarty et al., 2017) is another precision oncology knowledge base that allows searching and comparing drug response data from different TCGA cohorts. Although the ICGC web portal (Zhang et al., 2019) and Coral web application (Adelberger *et al.*, 2021) allow patient/gene subsetting of TCGA cohorts and provide survival and Venn diagram visualization of cohorts, they do not allow comparison of cohorts pregenerated by other research groups. These tools address the reuse needs of users. However, they only provide raw data without allowing the patient/gene subsetting of the statistical analyses for pan-cancer subcohort and associated gene discovery.

We built an interactive Shiny (Chang *et al.*, 2022) web application for the analysis and visualization of four data categories across 33 cancer types. Users can visualize the results of preprocessed analysis of Simple Nucleotide Variations (SNVs), Copy Number Variations (CNVs), differential gene expression in tumor versus normal samples, and clinical data of TCGA projects from National Cancer Institute (NCI) Genomic Data Commons (GDC) (Grossman *et al.*, 2016). Moreover, users can compare patient clusters determined using iClusterPlus R package (Mo and Shen, 2022) with expression-based survival risk groups (Zengin and Önal-Süzek, 2020, 2021), and curated subtypes such as immune subtypes (Thorsson *et al*, 2019), Triple Negative Breast Cancer (TNBC) subtypes (Lehmann *et al*, 2016), PAM50 subtypes (Berger *et al*, 2018) and Microsatellite Instability (MSI) related subgroups and several data type clusters from BiocOncoTK (Carey, 2022; Ding *et al*, 2018) and *curatedTCGAData* (Ramos *et al.*, 2020) R packages. Furthermore, users can create custom subcohorts based on genomic analyses and/or clinical data to subset data visualization. Users can also create gene sets for data type and/or pan-cancer comparisons. For each cancer, whenever available, sample types, survival risk groups (Low-risk / High-risk), and pre-computed or curated patient clusters can be used for filtering patients. The main novelty of our tool is allowing the users to generate custom patient sub-cohorts and/or gene sets using interactive graphical representations via clipboard functionality.

## 2 Methods

### 2.1 TCGA data

Publicly available hg38 data including SNV, CNV, Transcriptome Profiling, microRNA, Methylation, and clinical data of 33 cancer types from The Cancer Genome Atlas (TCGA) projects were downloaded on March 6, 2022 from NCI GDC (Grossman *et al.*, 2016) using TCGAbiolinks R package (Colaprico *et al.*, 2016).

### 2.2 Pre-computed Molecular Data Analysis

#### 2.2.1 SNV Analysis

Potential driver mutated genes with their roles as a tumor suppressor or oncogene were determined by SomInaClust R package (Van den Eynden *et al.*, 2015) using mutation annotation format (maf) file generated by mutect2 pipeline. With the “Somatic Driver Mutations” option, the user can see the significant mutated genes ranked by their Q-value. This option is only available for the “SNV Analysis” category.

#### 2.2.2 CNV Analysis

Significant recurrent copy number variations were identified by *GAIA* R package (Morganella *et al.*, 2011). NCBI IDs and Hugo Symbols of the genes on chromosomal regions with altered copy numbers were determined using *GenomicRanges* (Lawrence *et al.*, 2013) and *biomaRt* (Durinck *et al.*, 2009) R packages.

#### 2.2.3 Differential Gene Expression Analysis

Differentially expressed genes were determined using normalized HTseq counts, by limma-voom method with duplicate-correlation function from *edgeR* (Robinson *et al.*, 2010) and *limma* (Ritchie *et al.*, 2015) R packages. Ensembl IDs were converted to NCBI IDs and Hugo Symbols using the *biomaRt* package (Durinck *et al.*, 2009).

Two different analyses were performed using paired tumor samples against tumor-adjacent normal samples of patients with both sample types (Paired), or tumor samples of all patients against normal samples of patients who have both sample types (All) if it is available for a particular cancer.

#### 2.2.4 Pathway Enrichment

Pathway enrichment and visualization was performed for each analysis by *clusterProfiler* R package (Yu *et al.*, 2012).

### 2.3 Pre-computed Patient Clusters and Sample Subtypes

TCGAnalyzerR provides an interactive visual analysis of several patient cohorts: i) Survival Risk Groups: We provide low-risk or high-risk patient groups determined by expression-based gene signature analysis for Lung Adenocarcinoma (LUAD), Lung Squamous Cell Carcinoma (LUSC) and Colon Adenocarcinoma (COAD) (Zengin and Önal-Süzek, 2020, 2021), ii) iClusters: We clustered patients using their raw SNV, CNV, gene expression, miRNA expression and methylation data of tumor samples which have all types of data by iClusterBayes method (Mo *et al.*, 2018), iii) Curated subtypes (immune subtypes, TNBC subtypes, PAM50 subtypes) from original publications (Thorsson *et al*, 2019; Lehmann *et al*, 2016; Berger *et al*, 2018). For fifteen cancer types, previously published TCGA cohorts of the individual tumor types are retrieved from curatedTCGA R package curatedTCGAData R package (Ramos *et al.*, 2020). Patient clusters based on Microsatellite Instability (MSI) were compiled using BiocOncoTK (Carey, 2022, Ding *et al.*, 2018) and Immune clusters (Thorsson et al, 2019) were compiled for all 33 cancers. In total, 123 external patient cohorts are integrated into the web interface allowing efficient filtering and cross-comparative analysis of multiple subcohorts in parallel.

### 2.4 Survival Analysis

Kaplan-Meier (KM) survival analysis is performed by *readr* (Wickham *et al.*, 2022) and *survfit* (Therneau, 2022) R packages in real-time based on overall survival data of patients of interest for selected clinical features.

### 2.5 Visualization

TCGAnalyzeR front-end was implemented by javascript-based R packages with an interactive dashboard enabling users to select cancer types, data types, risk groups and patient cohorts using *heatmaply, g3viz* and *highcharter* R packages (Galili *et al.*, 2018; Guo *et al.*, 2019; Kunst, 2022). All visualizations are interactive and customizable by the user through the filtration options with “My genes” and/or “My patients” panels enabling to copy genes and/or patients of interest to the clipboard. <u>TCGAnalyzeR currently supports TSV for downloading tables; and high-resolution PNG format for downloading figures</u>.

## 3 Results

TCGAnalyzeR web application offers simple nucleotide (SNV) analysis as the first step. We present two data sets for SNV analysis: ‘Somatic Driver Mutations’ predicted by the SomInaClust R package and ‘All’ mutations from the original maf file without any analysis. Oncoplot in Figure 1 shows candidate driver genes with their percentage in tumor samples of Breast Invasive Carcinoma (BRCA) with annotations of patient iClusters, PAM50, TNBC and immune subtypes. iCluster #1 is highly correlated with the Basal and TNBC subtype. Wound-healing and IFNχ-dominant immune subtypes gather around iCluster #1. iCluster #2 is mostly correlated with Luminal A subtype and Inflammatory immune subtype. iCluster #3 seems to be a mixture of estrogen receptor positive Luminal A and Luminal B subtypes and heterogenous immune subtypes. Moreover, both iCluster #1 and iCluster #2 are not TNBC subtypes. On the other hand, iCluster #1 shows a highly different mutation pattern than other clusters. iCluster #1 together with basal and triple-negative subtypes have higher prevalence of TP53 mutations with very few mutations of PIK3CA, CDH1, GATA3, KMT2C, MAP3K1 genes. Moreover, mutations of TP53, CDH1 and GATA3 genes are mutually exclusive (Figure 1).

**Fig. 1.**
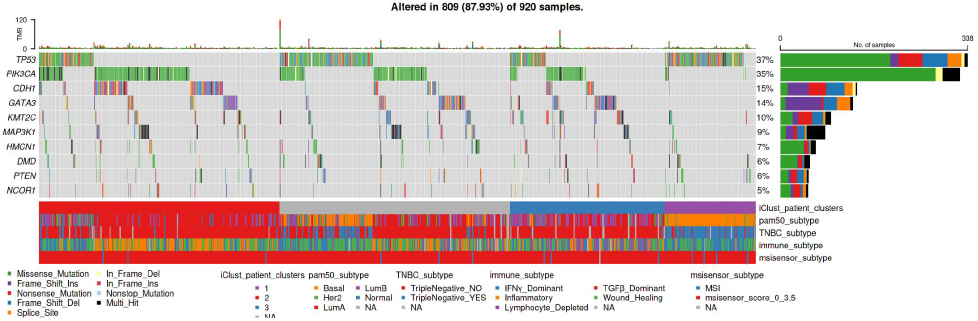
Oncoplot of candidate driver genes with patient clusters and sample subtypes. Top10 significant candidate driver genes with mutations determined by SomInaClust R t package. Bottom annotations show the sample subtypes curated from the literature and iClusters.

Pathway enrichment of candidate driver mutated genes is shown as a bar graph in Figure 2A. Significant pathways of driver genes are highly cancer-related pathways such as EGFR tyrosine kinase inhibitor resistance, PD-L1 and PD-1 pathway in cancer, prostate cancer, pancreatic cancer and chronic myeloid leukemia pathways. Pathway enrichment analysis also supplies a table showing KEGG IDs, with related genes and p/q-values (Figure 2B).

**Fig. 2.**
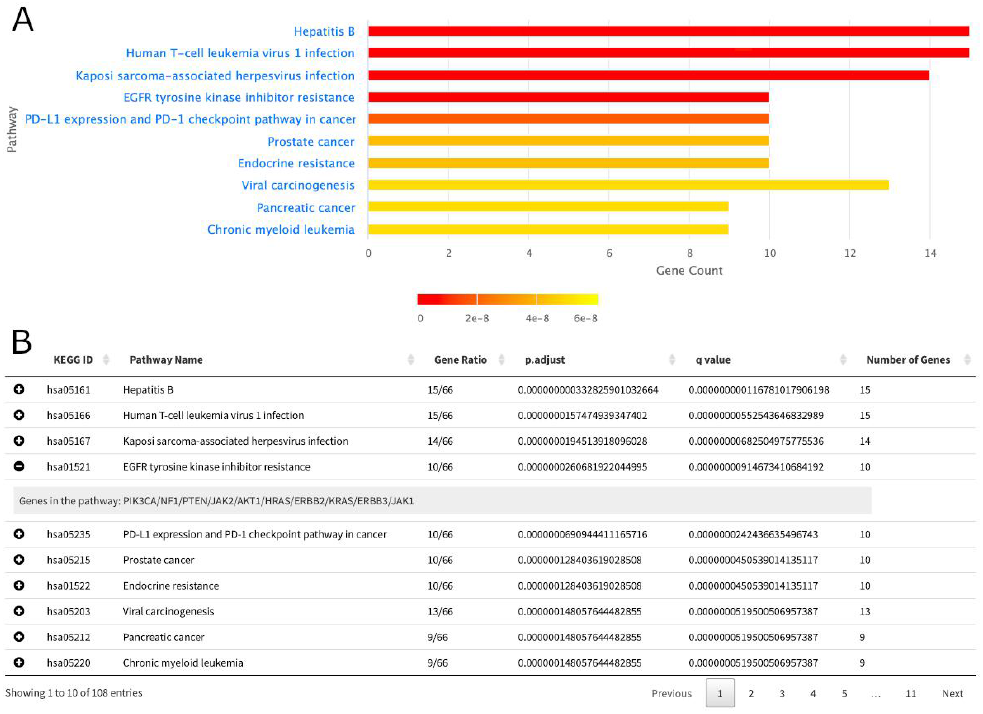
Pathway enrichment of candidate driver genes. A. Bar plot showing top 10 significant pathways of candidate driver genes determined by SomInaClust R package. B. Pathway enrichment table presenting KEGG ID, genes in significant pathways with adjusted p-value and q-value.

The “My genes” clipboard panel of TCGAnalyzeR allows modifying plots to show genes of interest. For example, Figure 3 shows the mutation pattern of Oncotype DX gene set together with clinical annotations. iCluster #2, Luminal A subtype and Her2 subtypes are related with ERBB2 (HER2) mutations. Besides, iCluster #1 have fewer mutations than the other two iClusters. Moreover, mutations of Oncotype DX genes are mostly mutually exclusive (Figure 3).

**Fig. 3.**
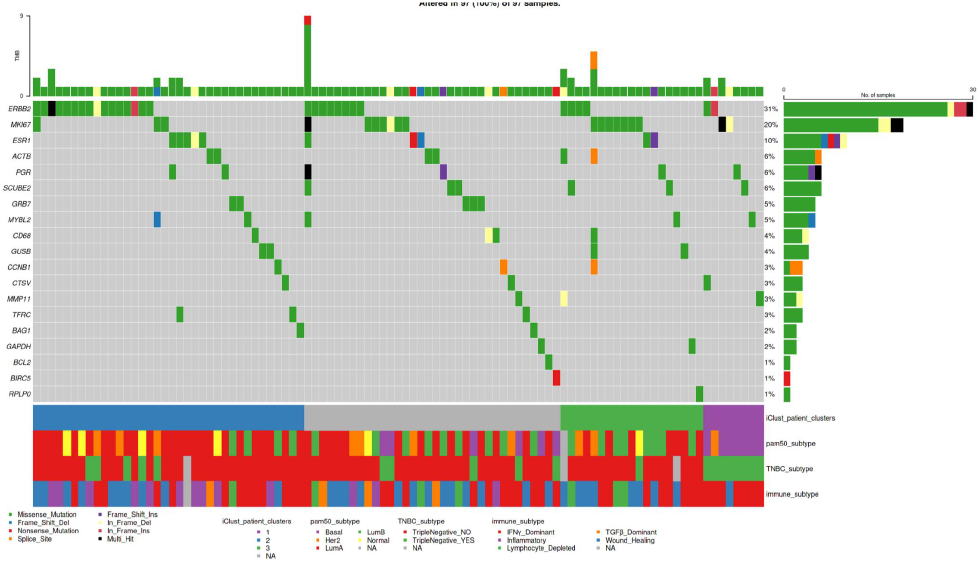
Oncoplot of Oncotype DX genes with patient clusters and sample subtypes. Mutations of Oncotype DX genes with annotations showing the patient iClusters and sample subtypes curated from the literature.

SomInaClust R package determines candidate driver mutated genes with their potential roles as a tumor suppressor (TSG) or oncogene (OG) with predicted scores (Van den Eynden *et al.*, 2015). Pyramid plot in Figure 4A summarizes OG score and TSG scores of candidate driver genes ranked by their analysis q-values. Some genes may have both OG score and TSG score over threshold, in that case, SomInaClust considers the COSMIC cancer gene census information. Only 1 (ERBB2) of 21 Oncotype DX genes were predicted as significant driver mutated genes. ERBB2 is predicted as an oncogene by SomInaClust as listed Dominant (OG) in COSMIC cancer gene census (Figure 4B).

**Fig. 4.**
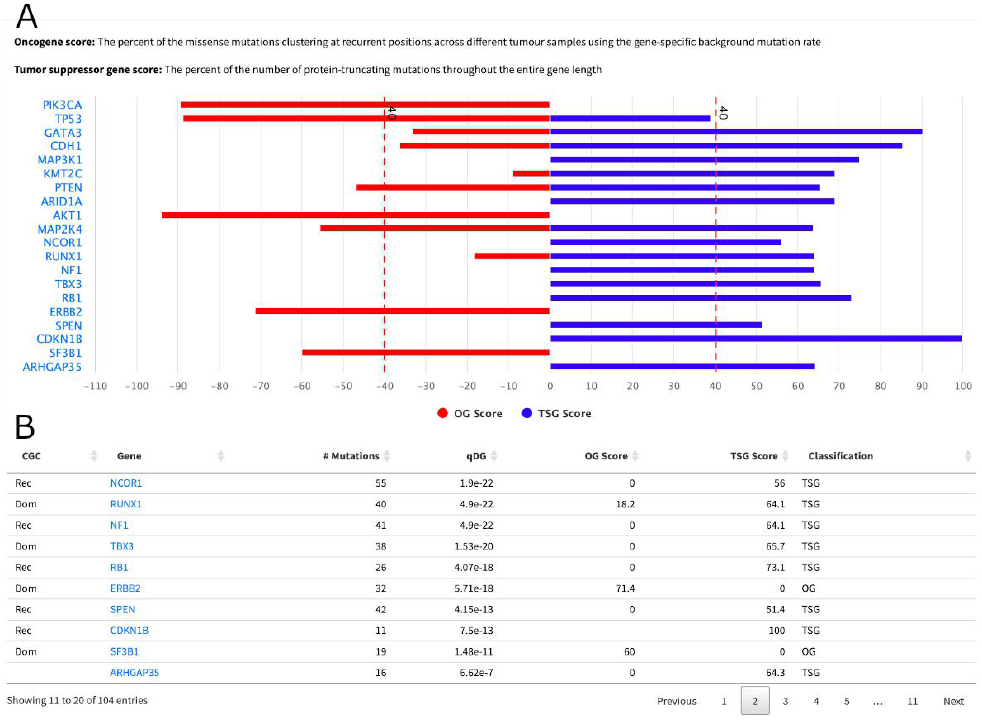
SomInaClust prediction of candidate driver genes. A. Pyramid plot showing Oncotype DX genes of which were predicted as candidate driver genes with calculated oncogene (OG) and tumor-suppressor (TSG) scores by SomInaClust R package. B. Detailed results of SomInaClust analysis with number of mutations, OG score, TSG score and q-value (qDG). CGC: COSMIC cancer gene census, Rec: Recessive (TSG), Dom: Dominant (OG).

Transcriptome analysis module provides differential expression analysis (DEA) of RNAseq data by comparing the expression of genes in primary tumor samples against adjacent normal samples. We present two results options for this analysis: ‘Paired’ as comparison of tumor samples against their own paired normal or ‘All’ as comparison of tumor samples against a normal sample subset of patients if such is available for the particular cancer. Volcano plot in Figure 5A summarizes the differential expression analysis of paired BRCA samples and Oncotype DX genes highlighted through the ‘My Genes’ panel. Table in Figure 5B presents the details of DEA with gene symbol, fold change (logFC) and p values of top 10 significantly differentially expressed genes ranked by p-value.

**Fig. 5.**
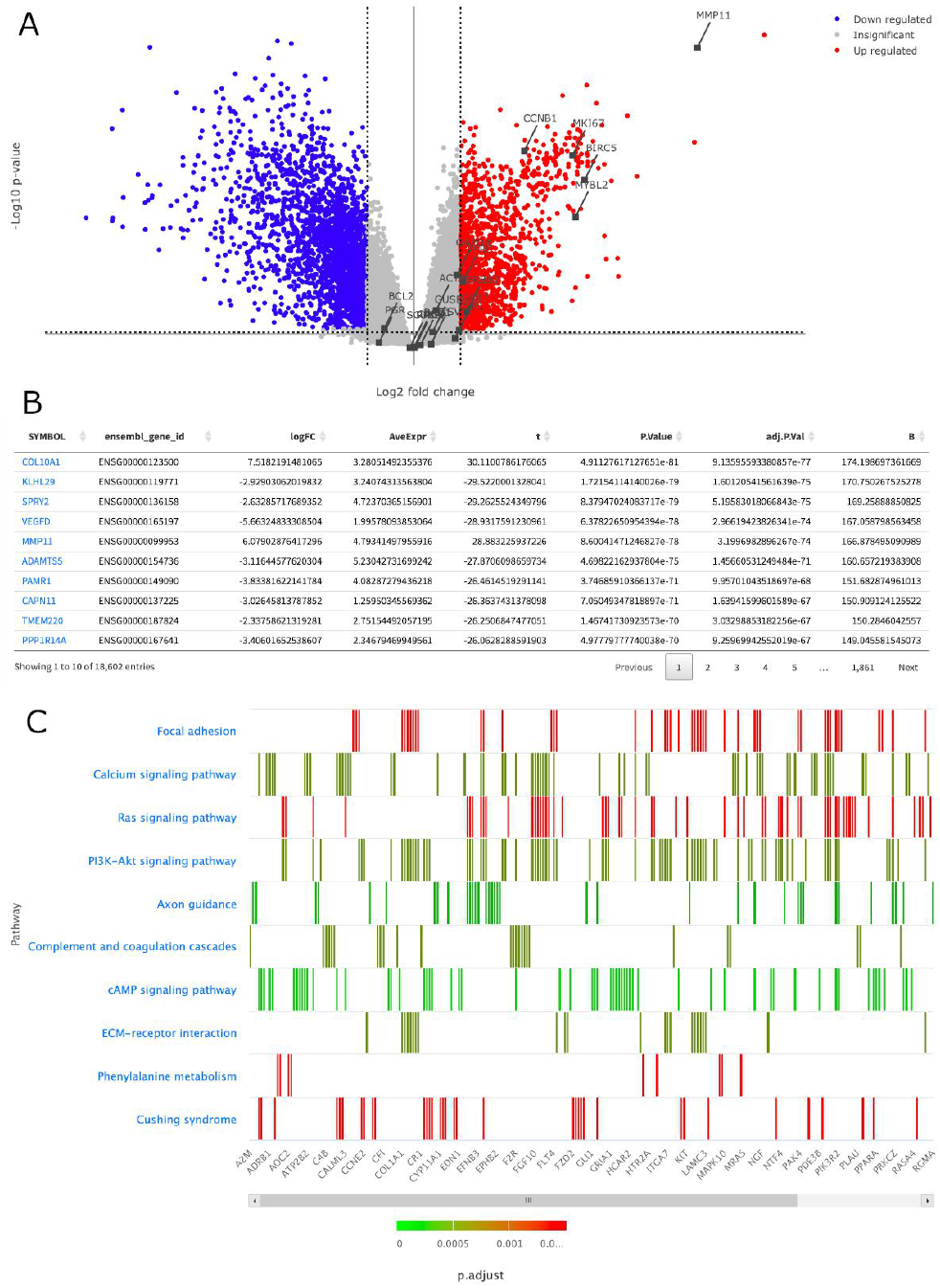
Differential expression of genes in tumor samples versus normal samples. A. Volcano plot showing up-regulated and down-regulated genes with −log10 conversion of p-values. Oncotype DX genes are highlighted on the graph. B. Differential expression results table presenting gene symbols, fold changes (logFC) and adjusted-p-values. C. Heatmap showing pathway enrichment of differentially expressed genes.

Pathway enrichment of differentially expressed genes showed that these genes play role in focal adhesion and ECM-receptor interaction which can be related with metastasis; Ras signaling, PI3K-Akt signaling, cAMP signaling and Phenylalanine metabolism pathways which are related with cell growth (Figure 5C). Genes related to these pathways can be observed with their p value color representation at heat plot of pathway enrichment analysis (Figure 5C).

Metastasis related gene MMP11, proliferation related genes BIRC5, MYBL2, MKI67 (Ki67), AURKA (STK15), CCNB1 and ERBB2 (HER2) from Oncotype DX gene set are highly up-regulated significantly in tumor samples of BRCA (Figure 5A). However hormone related genes (BAG1, BCL2, CD68, ESR1 (ER), GSTM1, PGR, SCUBE2) are not significantly differentially expressed among all tumor samples.

When we concentrate on the ERBB2 gene because it was predicted as a driver oncogene, we can visualize positions of mutations by the Lollipop plot in Figure 6A. Most of the mutations of ERBB2 gene are located on the kinase domain of HER2 (Figure 6A). These mutations are mostly missense on protein positions 755 (n=7), 767 (n=2), 769 (n=3), 777 (n=4), 797 (n=1), 842 (n=1), 939 (n=1) and in frame insertion on protein position 885 (n=1). Mutations on ERBB2 gene in tumor samples cause lower survival probability with 1.43 hazard ratio (not significant, p=0.084) (Figure 6B).

**Fig. 6.**
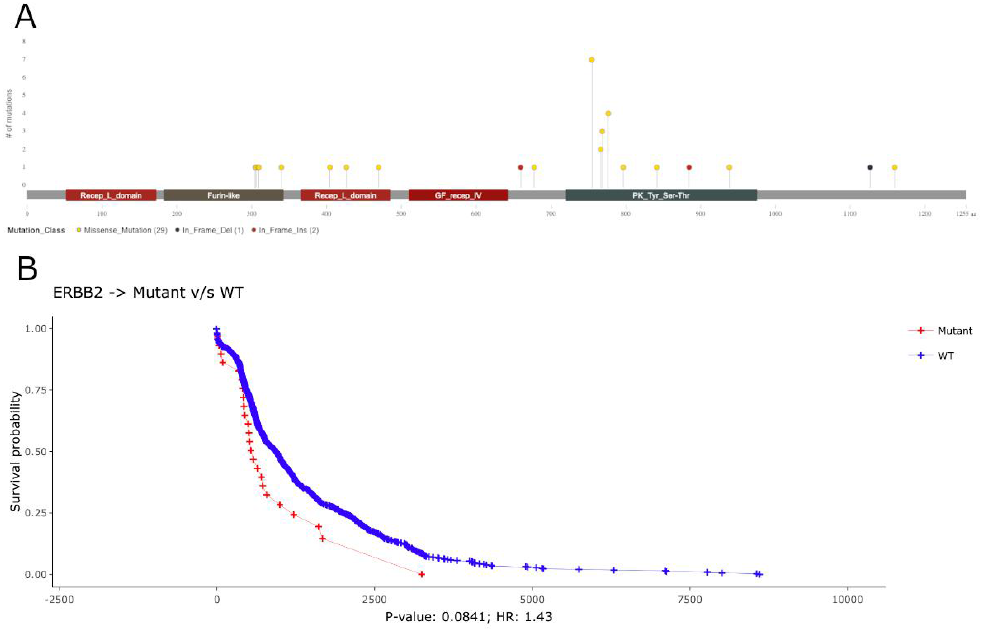
Detailed analysis of ERBB2 (HER2) mutations. A. Lollipop plot showing mutations of ERBB2 gene among tumor samples. B. Overall survival analysis of wild type versus mutated ERBB2 in tumor samples.

When we checked the expression levels of ERBB2 in tumor samples versus normal samples, from paired DEA, ERBB2 is expressed in tumor samples significantly higher than their adjacent normal samples (p=3.521E-10) (Figure 7A). However, patients with higher expression of ERBB2 have higher survival probability significantly (p=0.045) (Figure 7B).

**Fig. 7.**
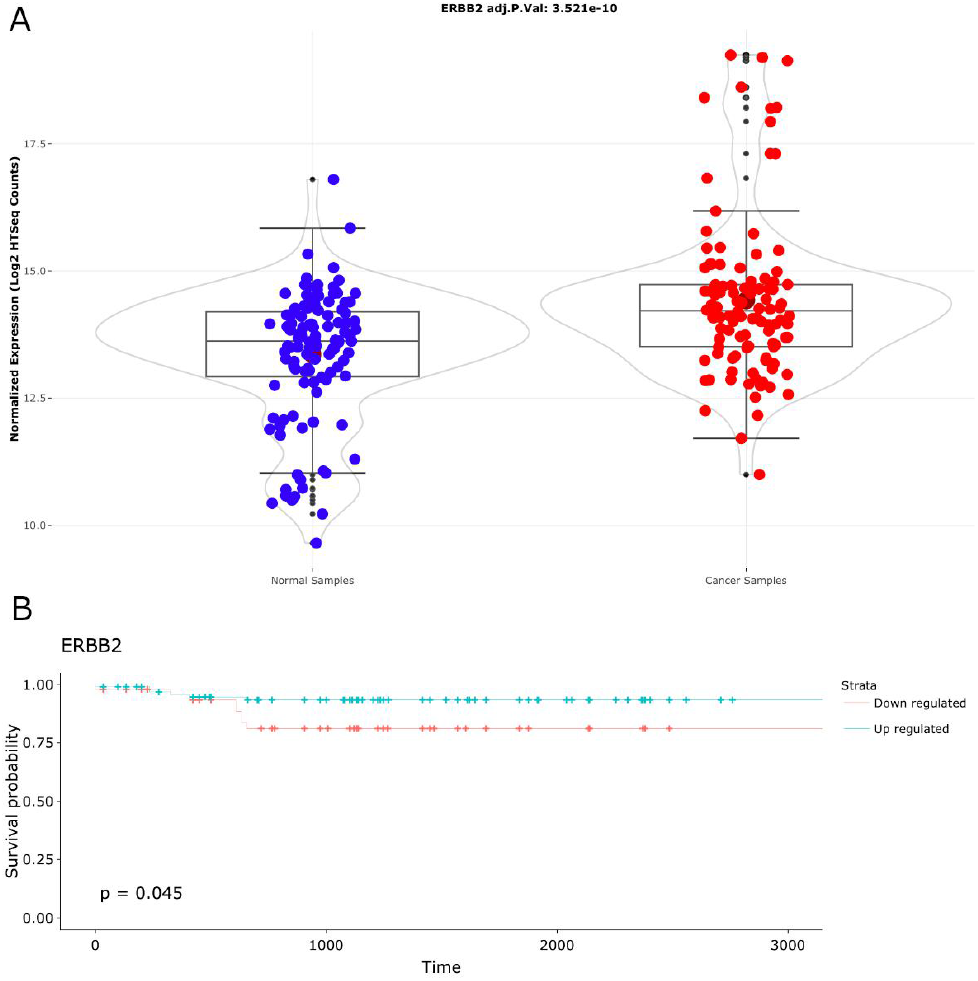
Detailed analysis of ERBB2 (HER2) expression. A. Violin plot presenting log2 transformed normalized mRNA expression of ERBB2 in normal and tumor samples with adjusted p-value. B. Overall survival analysis of expression levels of ERBB2 in tumor samples.

Clinical data analysis is composed of pie chart visualization and survival analysis of clinical features with curated patient clusters and sample subtypes. Figure 8 shows the visualization of iClusters, PAM50 subtypes and MSI-sensor subtypes. iClusters showed differential survival probability close to significance level (p=0.057) (Figure 8A), however PAM50 subtypes do not have differential survival probabilities. (p=0.68) (Figure 8B). Patients who have tumors with MSI have lower survival probability than patients with MSI stable tumors (p=0.0022) (Figure 8C).

**Fig. 8.**
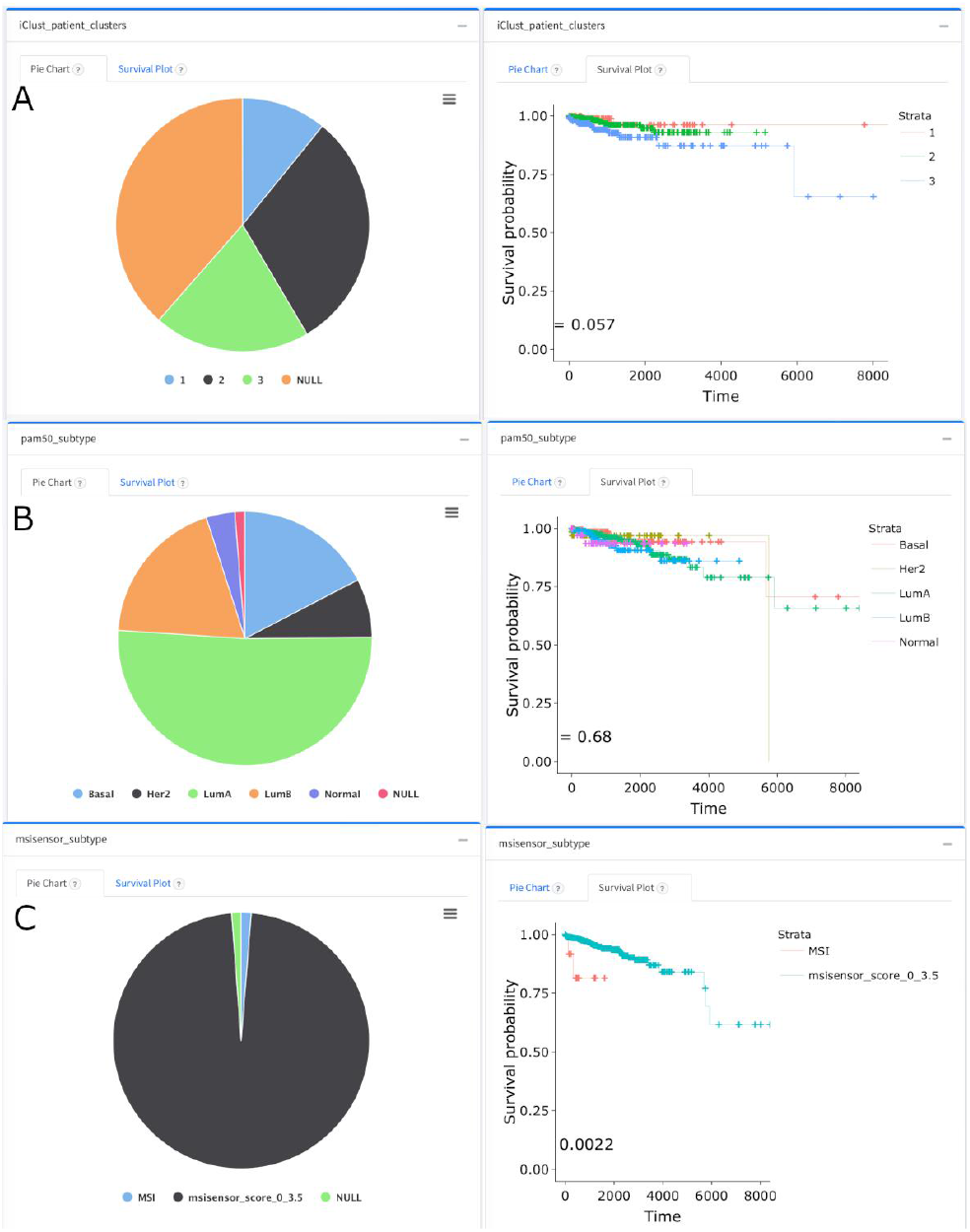
Pie charts and survival analysis of clinical features. A. Pie chart representation and survival analysis of iClusters. B. Pie chart representation and survival analysis of PAM50 subtypes. C. Pie chart representation and survival analysis of MSI subtypes.

Radial slices of the pie-chart are clickable, letting the user add the corresponding patient subsets to the ‘My Patients’ clipboard panel. Besides, users can customize a variety of plots such as survival plot, volcano plot, box plot, heatmaps, lollipop plot, and pie charts for discovering common molecular profiles for precision oncology. Each plot and data table are downloadable for use in articles.

## 4 Conclusion

Several web portals facilitating analysis on TCGA data have been developed and widely used such as Genomic Data Commons (GDC) data portal (Grossman, 2016), ICGC data portal (Zhang, 2019) and CPTAC data portal (Edwards, 2015). The cBioPortal is an open-access, open-source resource for interactive exploration of multidimensional cancer genomics data sets (Cerami, 2012; Gao, 2013) providing gene- centered query and visualization functions across multiple cancers. IntOGen is another similar framework for automated comprehensive knowledge extraction based on mutational data from sequenced tumor samples from TCGA patients (Francisco, 2020). However, we provide pre-performed SNV, CNV and differentially expression analyses with large sets of patient clusters and sample subtype information. We present signature based clustering using Generalized Linear Model for three cancer types (LUAD, LUSC and COAD). For all 33 cancer types immune and MSI-sensor scores of all patients are retrieved from their original publications. For the breast cancer (BRCA), PAM50 and TNBC patient cohorts are retrieved from their original publications. For fifteen cancer types, previously published TCGA cohorts of the individual tumor types are retrieved by curatedTCGAData R package (Ramos et al, 2020). By the time this manuscript is written iClusterPlus based patient cohorts are generated for fifteen cancers based on five data dimensions: miRNA, methylation, single nucleotide variation, transcriptome and copy number variation. A re-runnable parallel Linux pipeline is implemented enabling a scalable update of the pan-cancer data at the backend. We plan to generate iClusters for 33 cancer types, and integrate results of miRNA and methylation analyses, too. Since its initial inception to public in January 1st of 2022, TCGAnalyzeR has been regularly accessed by ~79 unique users/day.

TCGAnalyzeR provides a user-friendly web framework for integrative, large-scale analyses of genomic and clinical data of 33 cancer types from TCGA. TCGAnalyzeR web interface allows clinical oncologists and cancer researchers to create subcohorts and/or gene sets of interest to filter visualization of analyses. TCGAnalyzeR help page includes a demonstration of the app with the two use-cases of subcohort discovery and can be used as a manual.

## Funding

This work has been supported by Turkish National Institutes of Health (TÜSEB) [4583].

### Conflict of Interest

none declared.

## Notes

http://tcganalyzer.mu.edu.tr

### Competing Interest Statement

The authors have declared no competing interest.

### Summary of Updates

The title has changed. Few paragraphs are re-written.

